# A miRNA catalogue and ncRNA annotation of the short-living fish *Nothobranchius furzeri*

**DOI:** 10.1101/103697

**Authors:** Mario Baumgart, Emanuel Barth, Aurora Savino, Marco Groth, Philipp Koch, Andreas Petzold, Ivan Arisi, Matthias Platzer, Manja Marz, Alessandro Cellerino

**Author notes:** shared first authorship. shared last authorship; to whom correspondence should be addressed.

## Abstract

**Background:** The short-lived fish *Nothobranchius furzeri* is the shortest-lived vertebrate that can be cultured in captivity and was recently established as a model organism for aging research. Small non-coding RNAs, especially miRNAs, are implicated in age-dependent control of gene expression.

**Results:** Here, we present a comprehensive catalogue of miRNAs and several other non-coding RNA classes (ncRNAs) for *Nothobranchius furzeri*. Analyzing multiple small RNA-Seq libraries, we show most of these identified miRNAs are expressed in at least one of seven *Nothobranchius* species. Additionally, duplication and clustering of *N. furzeri* miRNAs was analyzed and compared to the four fish species *Danio rerio*, *Oryzias latipes*, *Gasterosteus aculeatus* and *Takifugu rubripes*. A peculiar characteristic of *N. furzeri* as compared to other teleosts was a duplication of the miR-29 cluster.

**Conclusion:** The completeness of the catalogue we provide is comparable to that of zebrafish. This catalogue represents a basis to investigate the role of miRNAs in aging and development in this species.

**Availability:** All supplementary material can be found online at http://www.rna.uni-jena.de/en/supplements/nothobranchius-furzeri-mirnome/.

## 1 BACKGROUND

The annual teleost *Nothobranchius furzeri* is a recent experimental animal model in biomedical research. In the wild, this fish inhabits ephemeral pools in semi-arid *bushveld* of Southern Mozambique. It has adapted to the seasonal drying of its natural environment by producing desiccation-resistant eggs, which can remain dormant in the dry mud for one and maybe more years by entering into diapause. Due to very short duration of the rainy season, the natural lifespan of these animals is limited to a few months. They represent the vertebrate species with the shortest captive lifespan of only 4-12 months and also the fastest maturation. In addition, they express a series of conserved aging markers and are amenable to genetic manipulations, making them an attractive model system for aging research (for a review see [8]).

Here, we provide a comprehensive microRNA (miRNA) catalogue for *N. furzeri*. MiRNAs are abundant non-coding RNAs between 18 and 24 nucleotides in length that are produced in a complex biosynthetic process starting from longer transcripts and are established as key players in the post-transcriptional regulation of gene expression. MiRNA genes can be hosted within an intron of a protein-coding gene (and their transcriptional regulation follows that of the hosting gene) or can arise from primary transcripts that are regulated independently of any protein-coding RNA. Several miRNAs are grouped in genomic clusters containing mostly two to six individual miRNAs with an intra-miRNA distance of <10 kb, which are co-transcribed, however unusually large clusters were also found in some species, like the miR-430 cluster in zebrafish consisting of 57 miRNAs [30, 46, 51]. The advantage of this accumulation is unclear, it could be possible that multiple loci are required to increase the copy-number and therefore the expression level of specific miRNAs in particular conditions, like miR-430 in the maternal-zygotic transition in zebrafish (*Danio rerio*) [12]. MiRNA genes are grouped into families based on sequence homology and can be defined as a collection of miRNAs that are derived from a common ancestor [13]. On the contrary, miRNA clusters may contain miRNAs belonging to different miRNA families, but are located in relative close proximity to each other. Both the evolutionary conservation of some miRNA families and the innovations leading to appearance of novel miRNAs are well-described. An expansion of miRNA inventory due to genome duplications in early vertebrates and in ancestral teleosts has been described [16].

MiRNAs bind target mRNAs, due to sequence complementarity in the seed region (nucleotides 2 - 7), mostly in the 3’ untranslated region, thereby silencing expression of the gene product via translational repression and/or transcript degradation. Up to now, several thousands of miRNAs have been predicted and identified in animals, plants and viruses and one single species can express more than one thousand miRNAs [14]. They frequently represent the central nodes of regulatory networks and may act as “rheostat” to provide stability and fine-tuning to gene expression networks [34, 39].

Before a sequence of the *N. furzeri* genome assembly became available [36], we could show by use of the *Danio rerio* reference from miRBase, that aging in *N. furzeri* brain displays evolutionary conserved miRNA regulation converging in a regulatory network centered on the antagonistic actions of oncogenic MYC and tumor-suppressor TP53 [2] and the expression of miR-15a and the miR-17/92 cluster is mainly localized in neurogenetic regions of the adult brain [7]. Two draft genome sequences for *N. furzeri* were recently produced [36, 50], in this paper, we now provide a comprehensive annotation of the *N. furzeri* miRNome based on a combination of Illumina based small RNA-Seq, different *in silico* prediction methods on the genome assembly and a final manual curation. Using the newly created miRNA reference, we analysed a large dataset of 162 small RNA-Seq libraries and report tissue-specific miRNA expression of conserved and non-conserved miRNAs in *N. furzeri*. We further used the *N. furzeri* reference to analyse the miRNA expression in other Nothobranchius species and one closely-related non-annual killifish species, which were previously used to analyse positive selection [36] to identify when in the evolutionary history of *N. furzeri* non-conserved miRNAs arose.

## RESULTS AND DISCUSSION

### Small RNA-Seq libraries

For this study 150 small RNA-Seq libraries of *N. furzeri* from different ages and tissues were sequenced, making a total of around 2.2 billion reads. In more detail, we had 75 libraries for both of the strains *N. furzeri MZM-0410* (in the following called MZM) and *N. furzeri GRZ*. We investigated the three tissues brain, liver and skin at five different ages in *N. furzeri GRZ* (5, 7, 10, 12, 14 weeks) and *N. furzeri MZM* (5, 12, 20, 27, 39 weeks) with five biological replicates each. The only exception are the *N. furzeri MZM* brain libraries where we have four biological replicates for each age, but an additional time point of 32 weeks with five replicates. Additionally, seven embryonic small RNA-Seq libraries of *N. furzeri* were sequenced (two of the strain *MZM-0403* and five of *MZM-0410*). After preprocessing all libraries a total of around 1.9 billion high-quality reads was used for further analysis (see Methods section and Supplement Tab. 1).

For each of the six killifish species *A. striatum, N. kadleci, N. rachovii, N. pienaari, N. kunthae* and *N. korthausae* we generated two biological replicates of small RNA-Seq libraries from the brain of mature animals. The average size per library was 24.5 million reads with a minimum of 16.8 million and a maximum of 36.1 million reads (for more details about the small RNA-Seq libraries see Supplement Tab. 1).

### Annotation of ncRNAs

We could identify more than 750 non-coding RNA (ncRNA) genes in the *N. furzeri* genome based on small RNA-Seq reads (see Tab. 1, Supplement Material 1 and Supplement Tab. 5). In line with other eukaryotes, we identified multiple copies of rRNA operons, tRNAs, several major spliceosomal RNAs, signal recognition particle (SRP) RNAs and one copy of a minor spliceosomal RNA set. Further housekeeping RNA genes, such as RNase P, RNase MRP, and the 7SK RNA are found as expected once in the entire genome. We annotated the widely distributed TPP riboswitch, capable of binding thiamine pyrophosphate and thereby regulating genes that are in charge of the thiamine balance. We could also identify more RNA elements directly involved in the regulation of gene expression (3 copies of *IRE1* – controlling iron responsive proteins, CAESAR – controlling tissue growth factor CTGF, DPB – controlling DNA polymerase *β*), localization of mRNAs (Vimentin3), DNA replication (four copies of the *Y RNA* gene, and Telomerase RNA TERC), or of unknown function (12 vault RNAs). Additionally, ncRNAs responsible for editing certain mRNAs have also been found (two copies of Antizyme FSE, one U1A polyadenylation inhibition element (PIE), 26 Potassium channel RNA editing signals (KRES), and six copies of *GABA3*). Two promising candidate long non-coding RNAs (lncRNAs), SPYR-IT1 and MEG8, were also included in the annotation, even though we were not able to identify all of their exons. Two vague candidates for XIST and MALAT can be viewed in the supplemental material. The MALAT-derived masc and men RNA gene was clearly detected in 42 copies throughout the genome of *N. furzeri*.

**Table 1.** Number of annotated ncRNAs.

| ncRNA class | No. | ncRNA class | No. |
| --- | --- | --- | --- |
| 18S rRNA | 3 | 7SK | 1 |
| 5.8S rRNA | 11 | Vault | 12 |
| 28S rRNA | 6 | TPP | 1 |
| 5S rRNA | 28 | IRE1 | 3 |
| tRNA | 570 | CAESAR | 1 |
| U1 | 6 | DPB | 1 |
| U2 | 8 | Vimentin3 | 1 |
| U4 | 8 | Antizyme FSE | 2 |
| U5 | 8 | U1A PIE | 1 |
| U6 | 6 | KRES | 26 |
| U11 | 1 | GABA3 | 6 |
| U12 | 1 | Y RNA | 4 |
| U4atac | 1 | TERC | 1 |
| U6atac | 1 | mascRNA-menRNA | 42 |
| RNase P | 1 | SPYR-IT1* | 1 |
| RNase MRP | 1 | MEG8* | 1 |
| SRP | 3 |  |  |

Besides housekeeping RNAs RNA elements controlling metabolism and protein editing were identified. \* lncRNA candidate.

## Mapping and miRNA prediction results

For the identification of putative miRNA genes we used five methods, each following a different prediction approach (`BLAST, CID-miRNA, Infernal, GoRAP, miRDeep*`) and `Blockbuster` as verification (see Fig. 2 for an example).

The five tools identified 71, 33, 407, 209, 490 miRNA candidates, respectively (Fig. 1 shows the variety and the overlap of the different tools). All predictions were merged and redundant loci were removed (for details see methods section). Of the remaining 788 candidate miRNAs 617 (78.3 %) showed expressions and were verified by `blockbuster` and manually investigated for miRNA-like profiles. By this 34 (4.3 %) candidates were removed, all predicted by miRDeep_*_. Candidates showing no expression in any of the sequenced small RNA-Seq libraries were still kept as putative miRNAs. In total we predict a final amount of 754 miRNAs in *N. furzeri* by union of these methods (see Supplementary Material 2).

**Fig. 1.**
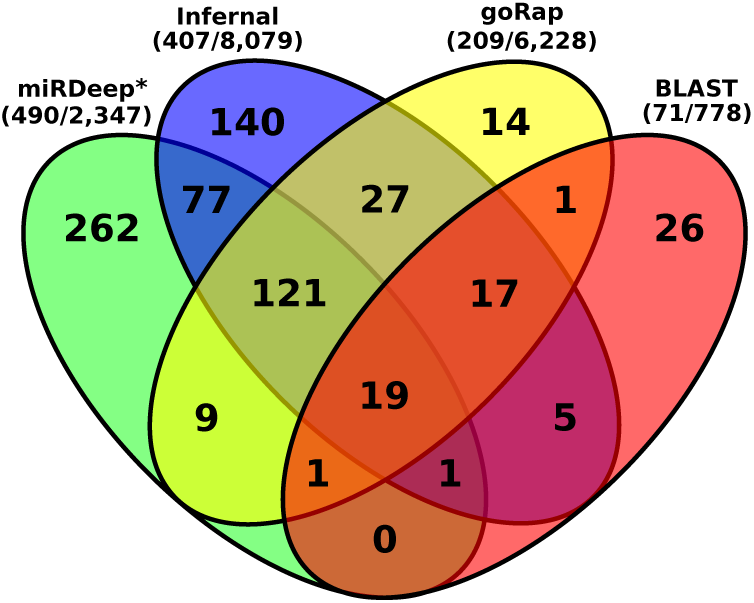
Venn diagram of predicted miRNA genes from four tools miRDeep_*_, Infernal, goRap and BLAST. Only 2 of the 33 candidates predicted by `CID-miRNA` overlapped with any of the other miRNA candidates. Nevertheless all 33 candidates were selected as miRNAs after manual inspectations. The total number of miRNA predictions after and before applying any filtering step are shown in brackets for each tool.

**Fig. 2.**
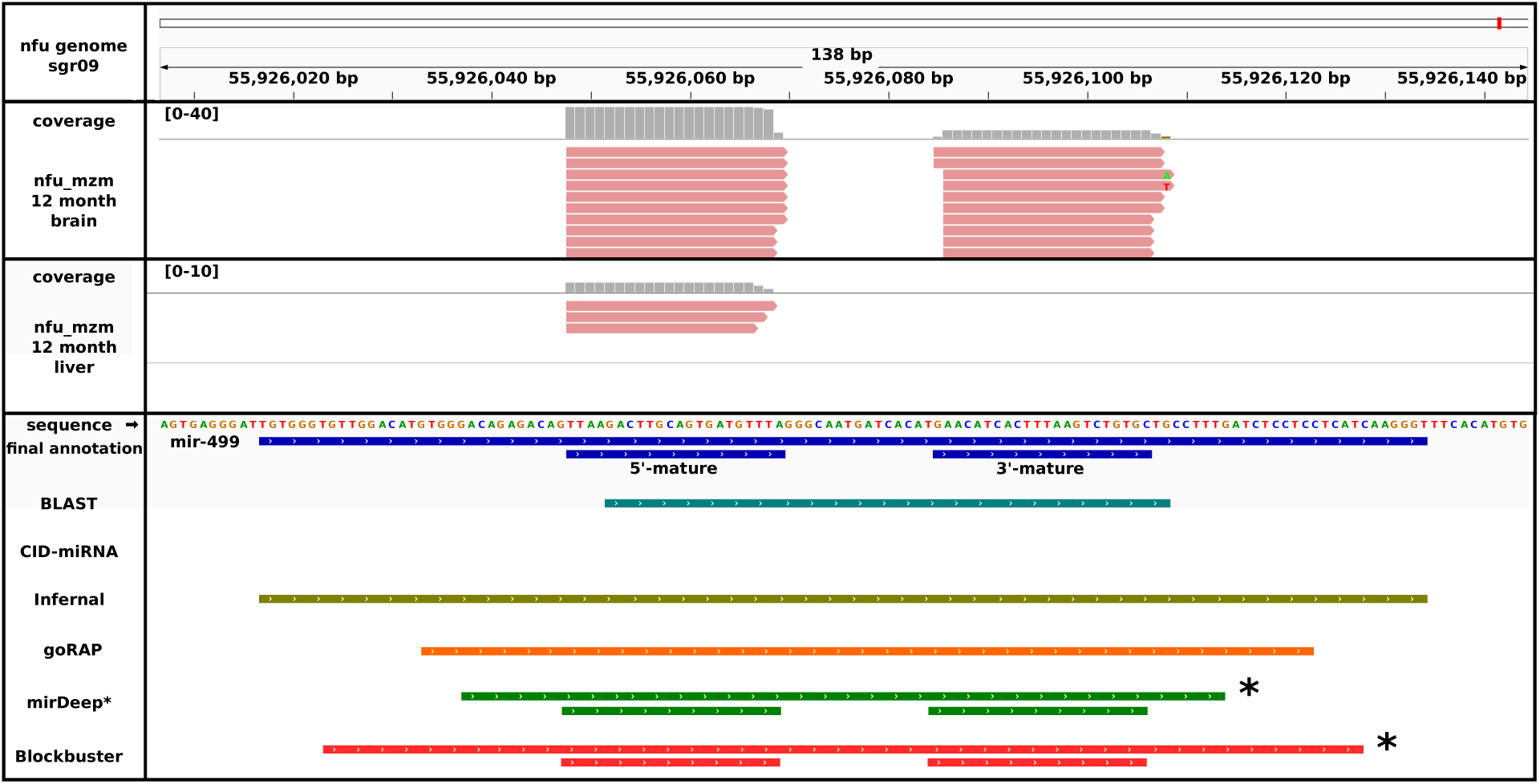
Annotation, expression profiles and prediction comparison for miR-499. We annotated the pre-miR-499 on sgr09, position 55,926,017–55,926,134 and the two mature miRNAs at 55,926,048–55,926,069 and 55,926,085–55,926,106. The six methods used for miRNA detection are displayed, CID-miRNA was not able to detect this miRNA. Tools working independent of the small RNA-Seq data (`BLAST` (cyan), `Infernal` (olive green) and `goRAP` (orange) vary in their annotation length. The latter two programs are based on covariance models identifying mostly the complete pre-miRNA. The remaining two programs miRDeep_*_ and `Blockbuster` are based on small RNA-Seq data (*) and therefore accurately annotate the mature miRNAs. MiR-499 is expressed weakly within *N. furzeri MZM* 12 month liver library and therefore could not be detected by miRDeep_*_ and `blockbuster`. In the *N. furzeri MZM* 12 month brain library miR-499 was expressed high enough to be detected by both programs.

Most of the small RNA-Seq reads (up to 88.81%) mapped onto the identified 754 miRNAs. Interestingly, the number of miRNA related reads varies broadly between the tissue samples (see Tab. 2). Possibly, this difference correlates with different regenerative capacities of these tissues. This suggests the existence of a class of ncRNAs, yet to be identified and annotated in *N. furzeri*, in line with a relatively high amount of reads mapping to unannotated locations in the liver samples. About half of the miRNA annotations are overlapping genes coding for proteins and are therefore intragenic. A minor fraction of reads (see Tab. 2) maps to other ncRNAs and proteins. Whereas 333 of the predicted miRNAs can be assigend to one of the known miRBase families, based on sequence identity, 421 miRNAs did not match any known family and can therefore be considered as novel or non-conserved miRNAs (for details see Table 3).

## Effects of tissue and age on global miRNA expression

We used principal component analysis (PCA) to visualize the effects of tissue type and age on global miRNA expression (see Fig. 3).

**Fig. 3.**
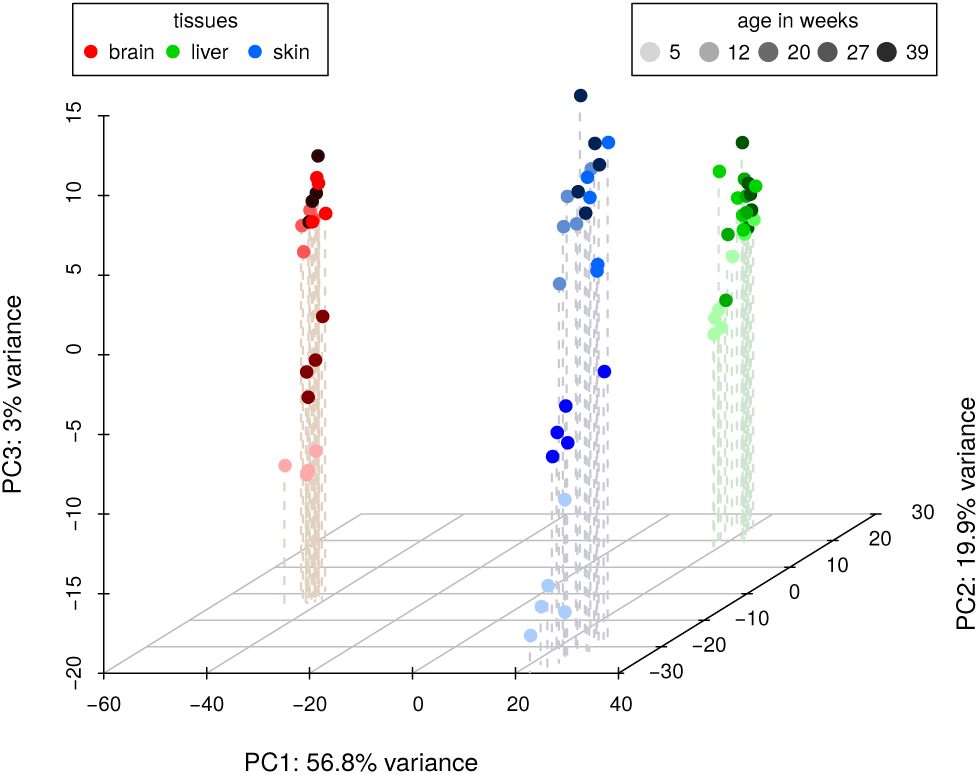
A three-dimensional PCA plot of the *N. furzeri MZM* small RNA-Seq libraries of all three tissues (brain – red, liver – green, blue – skin) and all investigated ages (from light to dark: 5, 12, 20, 27, 39 weeks). Whereas the samples cluster well according to their tissue belongings, a distinct separation regarding the ages can only be observed for the youngest samples in each tissue.

**Table 2.** Total number of genes known in *N. furzeri* and number of ncRNAs covered by small RNA-Seq reads and percental distribution of reads for brain, skin and liver of *N. furzeri*. See Supplement Tab. 1 for more details and Tab. 4 for details about the read libraries.

|  | Total | Brain |  | Skin |  | Liver |  |
| --- | --- | --- | --- | --- | --- | --- | --- |
| miRNA | 754 | 470 | 88.81 % | 438 | 75.01 % | 368 | 58.10 % |
| tRNA | 570 | 293 | 0.35 % | 361 | 1.33 % | 361 | 4.27 % |
| rRNA | 48 | 37 | 1.73 % | 43 | 3.61 % | 43 | 6.97 % |
| ncRNA | 250 | 136 | 0.09 % | 217 | 0.15 % | 204 | 0.41 % |
| protein* | 22,627 | 0 | 6.60 % | 0 | 4.59 % | 0 | 6.35 % |
| none |  |  | 2.42 % |  | 15.31 % |  | 23.90 % |
| Sum | 24,125 | 936 | 100 % | 1059 | 100 % | 976 | 100 % |

Asteriks indicates annotation of protein-coding genes from NCBI. ncRNA – All ncRNAs except miRNA, tRNA and rRNA; None – Reads mapping to non-annotated genomic areas.

**Table 3.**
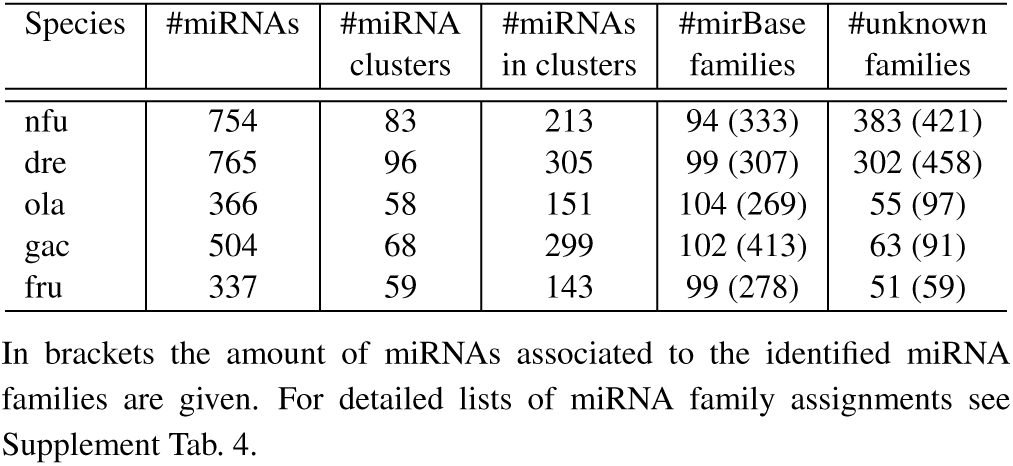
Amount of annotated miRNAs, identified miRNA clusters and the number of miRNAs in clusters, as well as known conserved and nonconserved miRNA families in *N. furzeri* (Nfu), *D. rerio (Dre), O. latipes (Ola), G. aculeatus (Gac) and T. rubripes (Fru).*

| Species | #miRNAs | #miRNA clusters | #miRNAs in clusters | #mirBase families | #unknown families |
| --- | --- | --- | --- | --- | --- |
| nfu | 754 | 83 | 213 | 94 (333) | 383 (421) |
| dre | 765 | 96 | 305 | 99 (307) | 302 (458) |
| ola | 366 | 58 | 151 | 104 (269) | 55 (97) |
| gac | 504 | 68 | 299 | 102 (413) | 63 (91) |
| fru | 337 | 59 | 143 | 99 (278) | 51 (59) |

A strong component of tissue-specific expression was detected and samples corresponding to different tissues clustered tightly and widely apart in the plane defined by the first two principal component axes (collectively accounting for 77 % of variance). Remarkably, the third principal component axis (3 % of variance explained) identifies an age-dependent component of miRNA expression, that is common to all three tissues with the youngest samples (5 weeks) clearly separated from the rest. A detailed analysis of age- and tissue-dependent miRNA expression including embryonic development will be part of a separate publication.

## MiRNA expression in closely related killifish

To compare and validate miRNA composition in *N. furzeri* we created for each of the six related killifish species two small RNA-Seq libraries (see Tab. 4). These libraries were mapped simultaneously on all available miRBase (release 21) sequences and our annotated miRNAs of *N. furzeri*, to observe *N. furzeri* miRNA-candidates expressed in other killifish and conserved miRNAs possibly missing in *N. furzeri*, but not in the closely related species. In total, 546 (93.7 %) of the 583 expressed and 17 (9.9 %) of the 171 non-expressed miRNA candidates in *N. furzeri* showed expression in at least one of the related killifish (Fig. 5 shows a miRNA not expressed in *N. furzeri* but in several of the other killifish). Of these expressed miRNAs 299 belong to the 421 non-conserved *N. furzeri* miRNA genes. To investigate whether miRNA sequences reflect known phylogenetic relationships, we concatenated the sequences of all expressed miRNAs and constructed a phylogenetic tree. This tree perfectly reflected the evolution of the Nothobranchius linage [10]. It is also interesting that the number of *N. furzeri* miRNAs expressed in other killifish species (indicated above the branch in Figure 4) is inversely correlated to the evolutionary distance, i.e. this number is higher for killifish the closer they are related to *N. furzeri*.

**Fig. 4.**
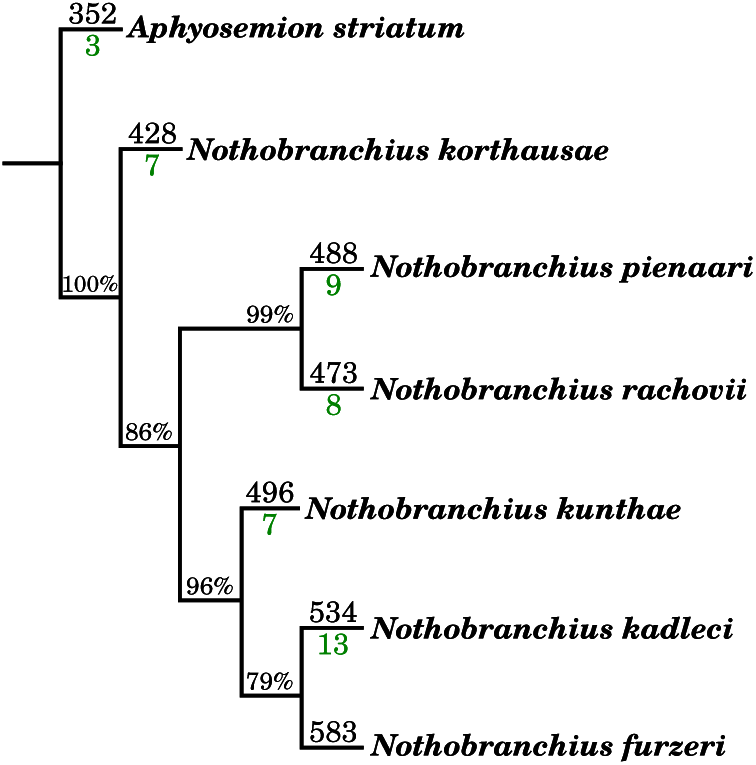
Killifish phylogeny based on the expressed miRNAs calculated via hierarchical clustering using the `R` package `pvclust` [41]. Bootstrap values are given as percentages at the corresponding branches. The amount of identified expressed miRNAs is given next to the species names. The numbers in green indicate the number of these expressed miRNAs, which were annotated but not expressed in any of the sequenced *N. furzeri* samples.

**Fig. 5.**
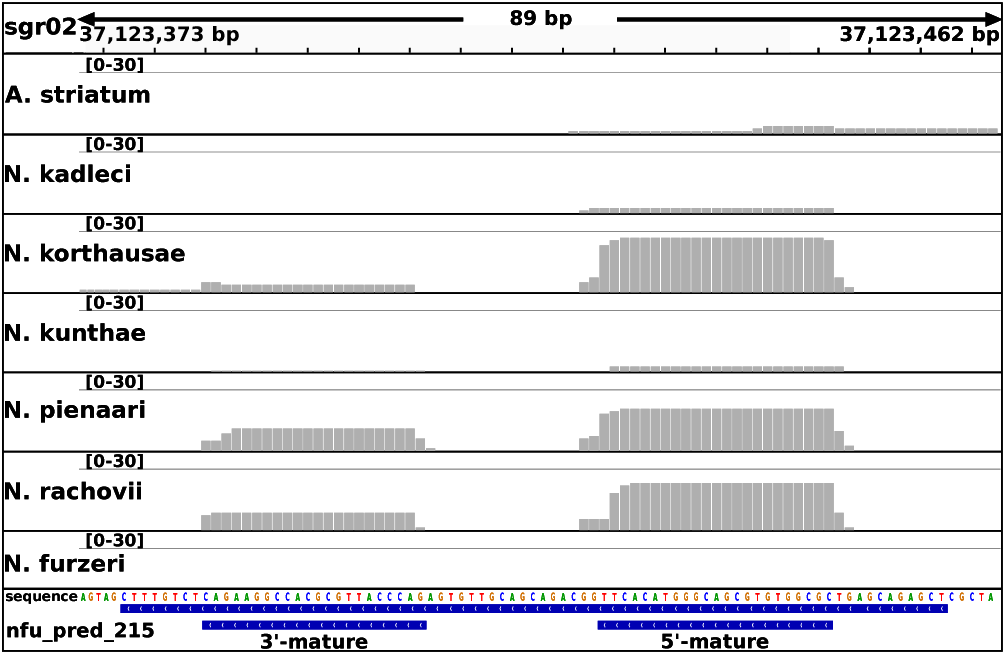
Expression profiles of the predicted miR-215. Grey bars indicate the amount of aligned reads and therefore coverage at the specific postions. Whereas no expression can be observed for this miRNA in *N. furzeri*, clear activation can be seen in *N. korthausae, N. pienaari and N. rachovii. A. striatum, N. kadleci* and *N. kunthae* show a weak expression for at least the 5’-mature variant of this miRNA.

**Table 4.**
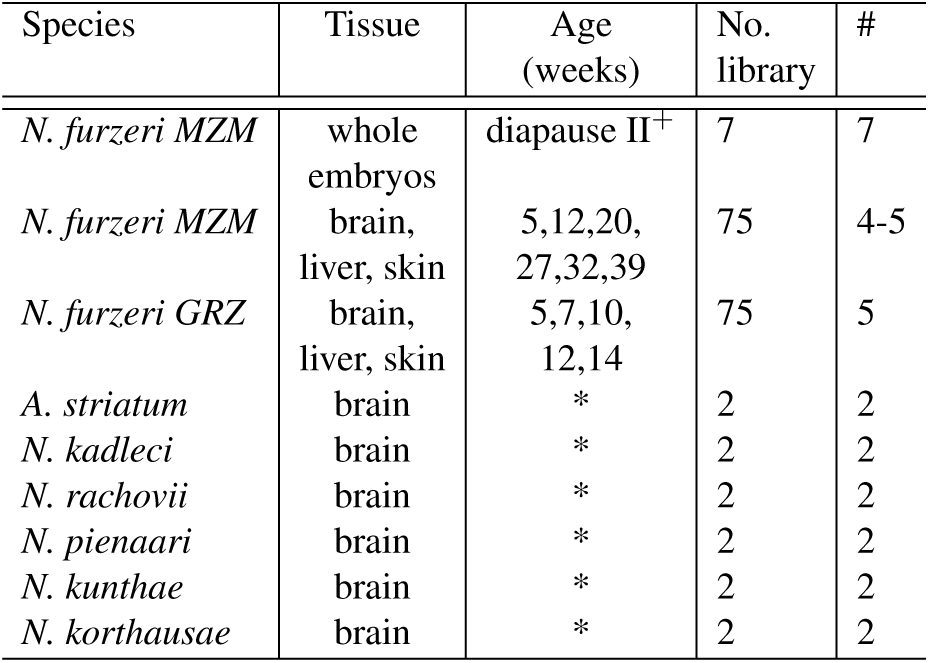
SmallRNA-Seq samples from Nothobranchius strains generated in this study. * – unknown; # – number of replicates; + – two weeks post-fertilization plus diapause

*A. striatum, N. korthausae, N. pienaari, N. rachovii, N. kunthae* and *N. kadleci* showed expression for 352, 428, 488, 473, 496 and 534 miRNAs, respectively. Most of these expressed miRNAs (> 89 %) are among the 333 conserved miRNAs of *N. furzeri* (see Supplement Tab. 3). The composition of expressed miRNAs from the six killifish varies only marginally. The *Nothobranchius* species (except *N. furzeri*) had in total 395 expressed miRNAs in common (of which 148 are non-conserved), and *A. striatum* expressed 324 of them (of which 116 are non-conserved). These 324 miRNAs represent a core of miRNAs from *Nothobranchiidae* whose origin predates the emergence of annualism in this clade.

## MiRNA clusters and gene duplications

MiRNAs are known to often occur in clusters [16]. We define a miRNA cluster to consist of at least two miRNAs with a maximum distance of 10 kb. Examining the localization and distances of the miRNA genes in the five fish species with assembled genomes, we identified 83, 96, 58, 68, and 59 different clusters in *N. furzeri*, *D. rerio, O. latipes, G. aculeatus and T. rubripes*, respectively (see Tab. 3, Fig. 6-A).

In all investigated fish species but *T. rupripe*s the largest cluster is the *mir-430* cluster (containing 7 to 55 miRNAs; see Fig. 6-C). This cluster is extremely divergent and evolving relatively quickly in each lineage. Not only the number of miR-430 copies within each cluster varies greatly, but also the number and organization of the members of this miRNA family. Whereas miR-430a and miR-430c can be found in all five fish species, miR-430b and miR-430d seem to occur only in *D. rerio and O. latipes* respectively.

Additionally, no structural similarities or shared repetition patterns can be observed for this miRNA cluster, which is an additional indication of the low purifying selection on this specific gene cluster. However, a clear duplication pattern can be observed for the miR-430 cluster in D. rerio (the order miR-430c/b/a is repeated with only few excpetions) and *N. furzeri* (the order miR-430c/a/a/c/a/a/a is repeated). For *O. latipes* and *G. aculeatus* the order of miR-430 variants appears to be more random and *T. rubripes* has too few copies to show any repeated pattern.

Fig. 6-B depicting the miR-17-92 cluster shows an example of the other extreme. In all five investigated fish species, two perfectly conserved clusters can be found. These represent a duplication of an ancestral cluster present in all vertebrates and the order of the different members is perfectly conserved. It is known that the miR-17-92 cluster is transcribed polycystronically and acts in oncogenic and tumor suppressor pathways [15, 43]. Furthermore, up to two smaller and lesser conserved clusters, containing at least two miRNAs of the miR-17 or miR-92 family were identified per fish species, similar to what is known for mammals. Having correctly identified this highly conserved cluster in *N. furzeri* is again good evidence for the high quality of its newly assembled genome and completeness of our miRNA catalogue.

**Fig. 6.**
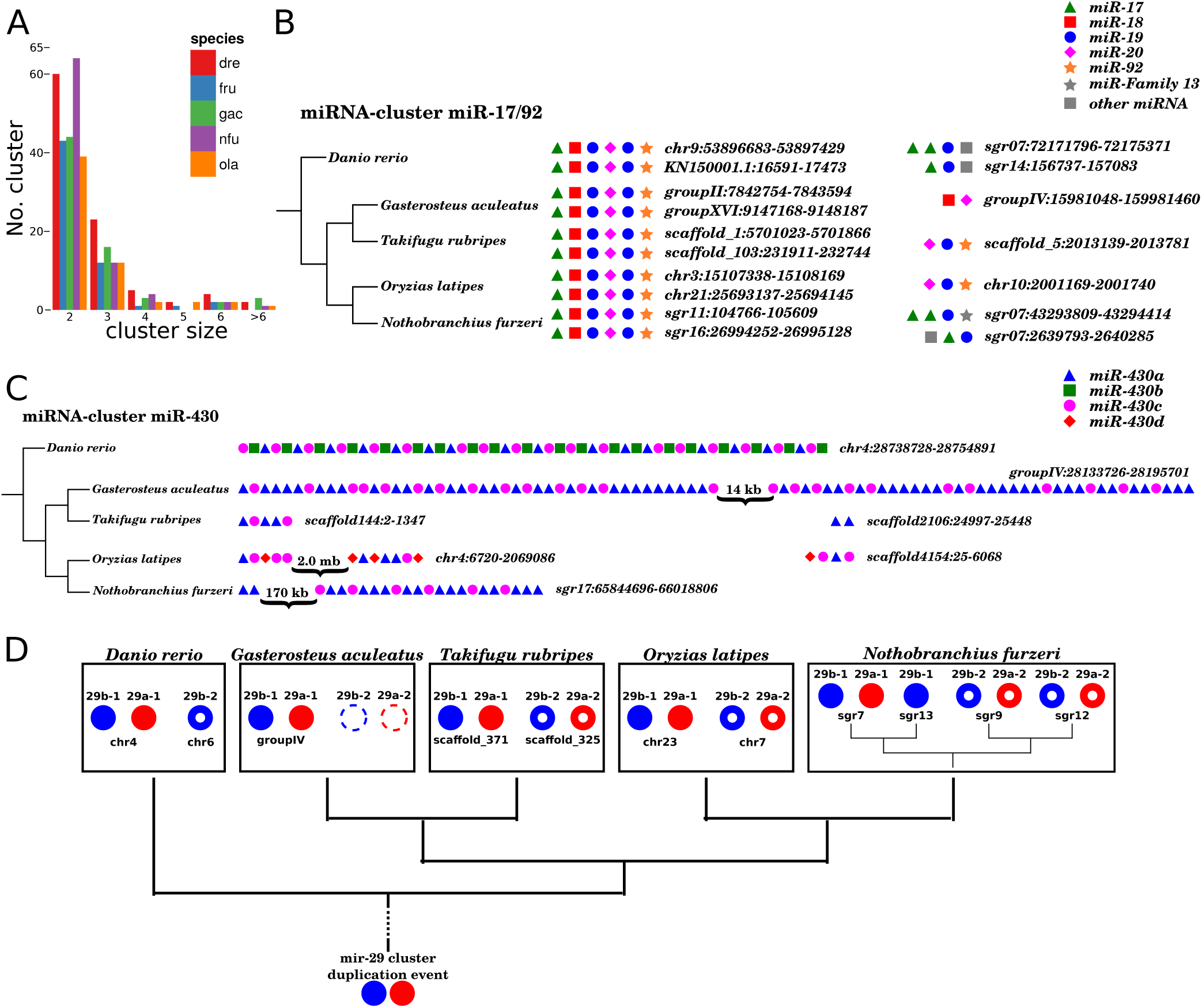
MiRNA cluster comparison between fish. (A) Amount of clusters and their respective sizes with a maximum distance of 10,000 bp between two miRNAs. (nfu - *Nothobranchius furzeri*, dre - *Danio rerio*, ola - *Oryzias latipes, gac* - *Gasterosteus aculeatus*, tru - *Takifugu rubripes*) (B) Structure comparison of the miR-17/92 cluster. Two highly conserved clusters could be identified for each species, as well as some smaller less conserved clusters, containing at least two miRNAs of the miR-17/92 cluster. (C) Structure comparison of the miR-430 cluster. No structural similarity between the different species can be observed. However, *D. rerio, D. aculeatus* and *N. furzeri* show some distinct but individual repeating pattern. Even though the gene variants miR-430b and miR-430d seem to be unique to *D. rerio* and *O. latipes*, they can be clearly distinguished, based on sequence alignments. (D) After the ancestral duplication event the mir-29 cluster is distinguished in the mir-29a/b-1 (filled red and blue dots) and the mir-29a/b-2 cluster (red and blue circles). Whereas for *D. rerio* the mir-29a-2 gene seems to be lost, we assume that for*G. aculeatus* the whole second mir-29 cluster (dashed circles) is only missing, because of the low quality genome sequencing and assembly. In *N. furzeri* we observe an additional copy for each of the two clusters, except that the mir-29a/b-1 pair was only partially duplicated or the second mir-29a-1 gene was lost again.

Another example for an evolutionary conserved miRNA cluster is the miR-29 cluster depicted in Fig. 6-D. Mir-29 family members are up-regulated during aging in a variety of different tissues including muscle, skin, brain and aorta [2, 11, 33, 40, 42, 49] and appear to be key regulators of age dependent gene expression [4, 37]. This cluster consists of miR-29a (which is identical to the mammalian miR-29c) and its variant miR-29b and is duplicated at leasts once. In some fish species an additional variant miR-29c is known, which is identical to the miR-29a in mammals with one nucleotide being different outside the seed region [29]. As from RFAM (version 12.1) and miRBase (release 21) miR-29 genes are mainly identified in vertebrates as well as one *Hemichordata* and one *Arthropoda*, so we can only speculate that the original cluster duplication event arose in the early metazoa lineage. In *O. latipes* and *T. rubripes* both miR-29 clusters are still present, whereas *D. rerio* seems to has lost one copy of the miR-29a gene. For *G. aculeatus* we were only able to identify one miR-29 cluster. However, since its genome assembly is incomplete we assume that the second cluster may not be lost but is missing in the current version of its miRNA annotation. Interestingly, in *N. furzeri* we identified an additional miR-29a/b pair and a fourth single copy of miR-29b. Assuming a complete genome assembly, different scenarios could explain this finding: (1) both original miR-29 clusters were individually duplicated once more and the fourth miR-29a gene was later lost, (2) one of the two clusters was duplicated as a whole, whereas in the other only miR-29b was copied, or (3) both original clusters were duplicated during the same event and again one of the miR-29a genes was later lost.

About the same amount of different miRBase miRNA families could be identified for all five fish species, despite their big differences in the number of identified miRNA genes. All miRNA genes not matching any known mirRBase family were clustered based on their sequence identity in order to estimate the amount of miRNA ‘families’ not covered by the miRBase database (see Tab. 3 and Supplement Tab. 4).

## CONCLUSION

This study involves a multitude of small RNA-Seq libraries from several tissues, ages, strains and embryos of *N. furzeri* and closely related species. The aim was the characterization of the *N. furzeri* miRNome and a detailed annotation in the recently published genome [36]. The inclusion of other killifish species allowed us to analyze the occurrence of novel miRNAs in the group of annual fish. Due to the fact, that we identified roughly the same number of miRNAs in *N. furzeri* as known in *D. rerio* and both fish species share almost equal amounts of miRBase families and unknown miRNA families, we assume that our miRNA catalogue is as complete as that of the model organism *D. rerio*.

## METHODS

### RNA extraction

Animal maintenance was performed as described [44, 45]. To avoid effects of circadian rhythms and feeding, animals were always sacrificed at 10 a.m. in fasted state. For tissue preparation, fish were euthanized with MS-222 and cooled on crushed ice. The protocols of animal maintenance and experiments were approved by the local authority in the State of Thuringia (Veterinaer- und Lebensmittelueberwachungsamt). Total RNA was extracted as described [2]. RNA quality and amount was determined using the Agilent Bioanalyzer 2100 and the RNA 6000 Nano Kit (Agilent Technologies).

### small RNA library preparation and sequencing

Library preparation and sequencing was done using *Illuminas* NGS platform [3]. One μg of total RNA was used for library preparation using *Illuminas* TruSeq small RNA sample preparation kit following the manufacturers instruction. The libraries were quantified on the Agilent DNA 1000 chip and subjected to sequencing-by-synthesis on an *Illumina* HiSeq2500 in high-output, 50 bp single-read mode. Sequencing chemistry v3 was used. Read data were extracted in FastQ format using the Illumina supported tool `bcl2fastq` (v1.8.3 and v1.8.4). The only exceptions were three of the *N. furzeri* embryo samples, which were sequenced on an *Illumina* HiSeq2000 in 50 bp single-read mode and where read data was extracted in FastQ format using the tool `CASAVA` (v1.8.2). Sequencing resulted in around 4 – 50 million reads per sample with pooling eight samples per lane.

In total 169 small RNA-Seq libraries from seven different killifish species were created. 157 of them were obtained from *N. furzeri* strains *GRZ* and *MZM-0410* at several ages from the three tissues brain, liver and skin. The remaining RNA-Seq libraries obtained from *Aphyosemion striatum, N. kadleci, N. rachovii, N. pienaari, N. kunthae and N. korthausae* were used to identify expression patterns at predicted miRNA locations in *N. furzeri* and miRBase pre-mature miRNA sequences. For details see Tab. 4, Supplement Tab. 1 and Supplement Tab. 2.

### Validation of mature miRNA expression by quantitative real-time PCR

For validation of mature miRNAs expression quantitative real-time PCR was performed an published in [2]. The used primers were Qiagen miScript primer: tni-miR-15a (cat# MSC0001523), tni-miR-101a (MSC0001279), tni-miR-101b (MSC0001283), dre-miR-145 (MSC0001529), has-miR-29c-1 (which is 100 % identical to dre-miR-29a, MS00003269), hsa-let-7a-5p (MS00031220), hsa-miR-124a-1 (MS00006622), has-miR-1-2 (MS00008358), ola-miR-21 (MS00049546), ola-miR-183-5p (MS00049392), and from cluster dre-miR-17a/18a/19a (MSC0002184–MSC0002186), and dre-miR-20a (MSC0002183).

### Small RNA-Seq library processing and mapping

In-house scripts were used to cut the RA3 adapter of the TruSeq small RNA preparation kit (5’-TGGAATTCTCGGGTGCCAAGG) from the reads. Additionally, `PRINSEQ` (v0.20.3) [38] was used to trim the reads from both sides, in order that the read bases had an minimum quality of 20 and reads were at least 15 bases long. Mapping onto the *N. furzeri* genome was performed with `segemehl` (v0.2.0) [18] using the -H 1 option, allowing single reads to be mapped to multiple best fitting locations. Visualization of mapped reads was done using `IGV` (v2.0.34) [47]. Since Bowtie (v1.0.0) [28] is the built-in method in miRDeep_*_ for mapping it was also used for the genomes of *N. furzeri*, *D. rerio, O. latipes* and *T. rubripes*.

### Genomes and annotations

The recently published high-quality draft genome assembly and annotation of *N. furzeri* and the small RNA-Seq libraries described above were used for mapping as well as for miRNA and other ncRNA predictions and annotations [36]. Additionally, these RNA-Seq libraries were also mapped on the following fish genomes: *Danio rerio* (GRCz10) [19], *Oryzias latipes* (HdrR) [22] and *Takifugu rubripes* (FUGU5) [20]. For annotation comparison, the latest complete genomic information of those three fish and of *Gasterosteus aculeatus* (BROAD S1) [20] were downloaded from the ensembl database [9].

### ncRNA and miRNA annotation

Non-coding RNAs in general were annotated with `GoRAP 2.0`, which is based on the RFAM database currently holding 2,450 ncRNA families (v12.0) [31].

For an initial prediction of candidate miRNAs a combination of five tools was used, each of them following a different annotation strategy: miRDeep_*_ (v32) [1], `Infernal` (v1.1) [32], `BLAST` (v2.2.30) [6], `GoRAP` (v2.0, unpublished) and `CID-miRNA` (version from April 2015) [48]. A detailed description of the individual searches can be found below.

All results were merged and putative miRNAs overlapping with genes of the recently published *N. furzeri* annotation were removed. Expression profiles of the remaining non-redundant candidate miRNA genes were analyzed automatically using `blockbuster` (v1) [26] and in-house scripts in order to mark candidates that did not exhibit a typical miRNA expression profil (according to [27, 21]). All candidates were additionally manually examined and filtered, leading to the final set of miRNA predicitions.

*MiRDeep* search* Mappings of 39 MZM brain, 15 GRZ brain, 25 GRZ liver, 28 MZM liver, 3 MZM skin and 7 MZM embryo small RNA-Seq libraries were used on four different fish genomes (*N. furzeri*, *D. rerio, O. latipes, T. rubripes*) as input for miRDeep_*_ (for a detailed list of used libraries see Supplement Tab. 1). Predictions from all 117 mappings were pooled together, in order to obtain a comprehensive representation of the miRDeep_*_ results. To each predicted miRNA hairpin sequence, we assigned the average of the miRDeep_*_ score computed across the multiple samples were the sequence was found.

The merged non-redundant list of identified miRNA sequences was re-mapped with `BLAT` [23] on the *N. furzeri* genome, and only gap-free alignments were accepted. These loci underwent further filtering steps: (i) A hairpin sequence was considered reliable if it showed a `BLAT` hit (one mismatch allowed) in miRBase (release 20) [24] or a miRDeep* score >= 7 and (ii) overlapping hairpin loci (i.e., within 100 nt) were discarded and the sequence with the highest score was kept. Predictions where no hits in miRBase could be obtained were further analyzed based on their secondary structure. Therefore, corresponding sequences were extended by 50 nt on either side and were compared with Rfam using `Infernal`. All predicted loci that had a significant hit to a known miRNA secondary structure or no hit at all were kept, while loci hitting other ncRNAs were discarded.

*Infernal search* For the `Infernal` search on the *N. furzeri* genome 155 hand curated pre-miRNA covariance models were used as input [5, 17] and only significant hits with a p-value of *p* < 0.005 were kept.

*BLAST search* In order to identify candidates from the most conserved miRNA families, `blastn` was used with all mature and pre-mature miRNA sequences available on miRBase (release 21) [25]. Only non-redundant hits were kept if they spanned the complete sequences of their corresponding input miRNAs to at least 90 % with no gaps allowed. To further reduce false positive hits a stringent cutoff of *p* < 10^−7^ was chosen.

*CID-miRNA* search Being based on a stochastic context free grammar model to identify possible pre-miRNAs, `CID-miRNA` follows a similar approach as `Infernals` covariance models. The *N. furzeri* genome was given as input with the following thresholds: putative miRNAs have a length between 60 bp and 120 bp and the grammar and structural cutoff were set to the recommended values of -0.609999 and 23, respectively.

## DECLARATIONS

### Ethics approval and consent to participate

Not applicable.

### Consent for publication

Not applicable.

### Availability of data and material

Supplementary material can be found online at. Data were deposited in GEO with the accession number GSE92854. The presented miRNome annotation is accessible and downloadable via the NFINgb Nothobranchius furzeri Genome Browser (http://nfingb.leibniz-fli.de/).

## Authors contribution

MB, MP and AC conceived and designed the study; MB performed the experiments; MG performed the RNA-Seq; IA, AP, EB, AS and AM analyzed and interpreted the data; AC, MB, EB and MM wrote the manuscript with contributions from all other authors.

## Acknowledgement

We thank Sabine Matz, Ivonne Heinze and Ivonne Goerlich for excellent technical assistance.

